# Sex-specific transcriptomic and epitranscriptomic signatures of PTSD-like fear acquisition

**DOI:** 10.1101/2021.11.25.468910

**Authors:** Andre Martins Reis, Jillian Hammond, Igor Stevanovski, Jonathon C Arnold, Iain S. McGregor, Ira Deveson, Anand Gururajan

**Author notes:** Corresponding Author: Anand Gururajan. Co-last authors (Ira Deveson, Anand Gururajan).

## Abstract

Our understanding of the molecular pathology of posttraumatic stress disorder (PTSD) is rapidly evolving and is being driven by advances in sequencing techniques. Conventional short-read RNA sequencing (RNA-seq) is a central tool in transcriptomics research that enables unbiased gene expression profiling. With the recent emergence of Oxford Nanopore direct RNA-seq (dRNA-seq), it is now also possible to interrogate diverse RNA modifications, collectively known as the ‘epitranscriptome’. Here, we present our analyses of the male and female mouse amygdala transcriptome and epitranscriptome, obtained using parallel Illumina RNA-seq and Oxford Nanopore dRNA-seq, associated with the acquisition of PTSD-like fear induced by Pavlovian cued-fear conditioning. We report significant sex-specific differences in the amygdala transcriptional response during fear acquisition, and a range of shared and dimorphic epitranscriptomic signatures. Differential RNA modifications are enriched among mRNA transcripts associated with neurotransmitter regulation and mitochondrial function, many of which have been previously implicated in PTSD. Very few differentially modified transcripts are also differentially expressed, suggesting an influential, expression-independent role for epitranscriptional regulation in PTSD-like fear-acquisition. Overall, our application of conventional and newly developed methods provides a platform for future work that will lead to new insights into and therapeutics for PTSD.

## Main text

The psychology of posttraumatic stress disorder (PTSD) is rooted in the acquisition of memories associated with specific traumatic events^1^. Neuromolecular changes in brain structures such as the amygdala, which play critical roles in learning and memory, have been implicated in the pathophysiology of PTSD^2^. Several studies have identified the impact of PTSD trauma on the epigenome^3, 4^. However, over the last decade, there is increasing evidence implicating RNA modifications, collectively known as the ‘epitranscriptome’, in brain function and behaviour^5-8^. These modifications can impact RNA stability, localisation and regulate translation in a dynamic and reversible manner. As such, they have the potential to rapidly fine-tune responses to external stimuli and may underpin aspects of fear learning and memory.

Here we report the results of our study that focused on capturing changes in the amygdala transcriptome and epitranscriptome during the acquisition of the conditioned fear response. We leveraged parallel short-read RNA-seq (Illumina NextSeq) and long-read direct RNA-seq (dRNA-seq; Oxford Nanopore Technology PromethION) to profile gene expression and RNA modifications, respectively, in a mouse model for aspects of PTSD.

Our PTSD mouse model was developed using a modified auditory, cued-fear conditioning protocol developed by Siegmund and Wotjak^9^ and has been validated for its face, construct and predictive validities^10-14^. Epidemiological evidence suggests that females are at higher risk (2:1) of developing PTSD following trauma^15^. However, to our knowledge, few PTSD-stress paradigms, including the one we have employed here, have been applied in females^16^. Accordingly, for our first experiment, we investigated if cued-fear memory was observable in adult female and male mice 24h after fear conditioning (**Figure 1a**). We observed a significant increase in freezing behaviour in both male and female conditioned (shocked) mice compared to non-conditioned (non-shocked) controls, with females freezing significantly more than males (**Figure 1b**). Our findings were consistent with one other study which also reported greater cued fear expression in females than males, suggestive of sex-differences in discriminative ability to the cue^17^.

**Figure 1.**
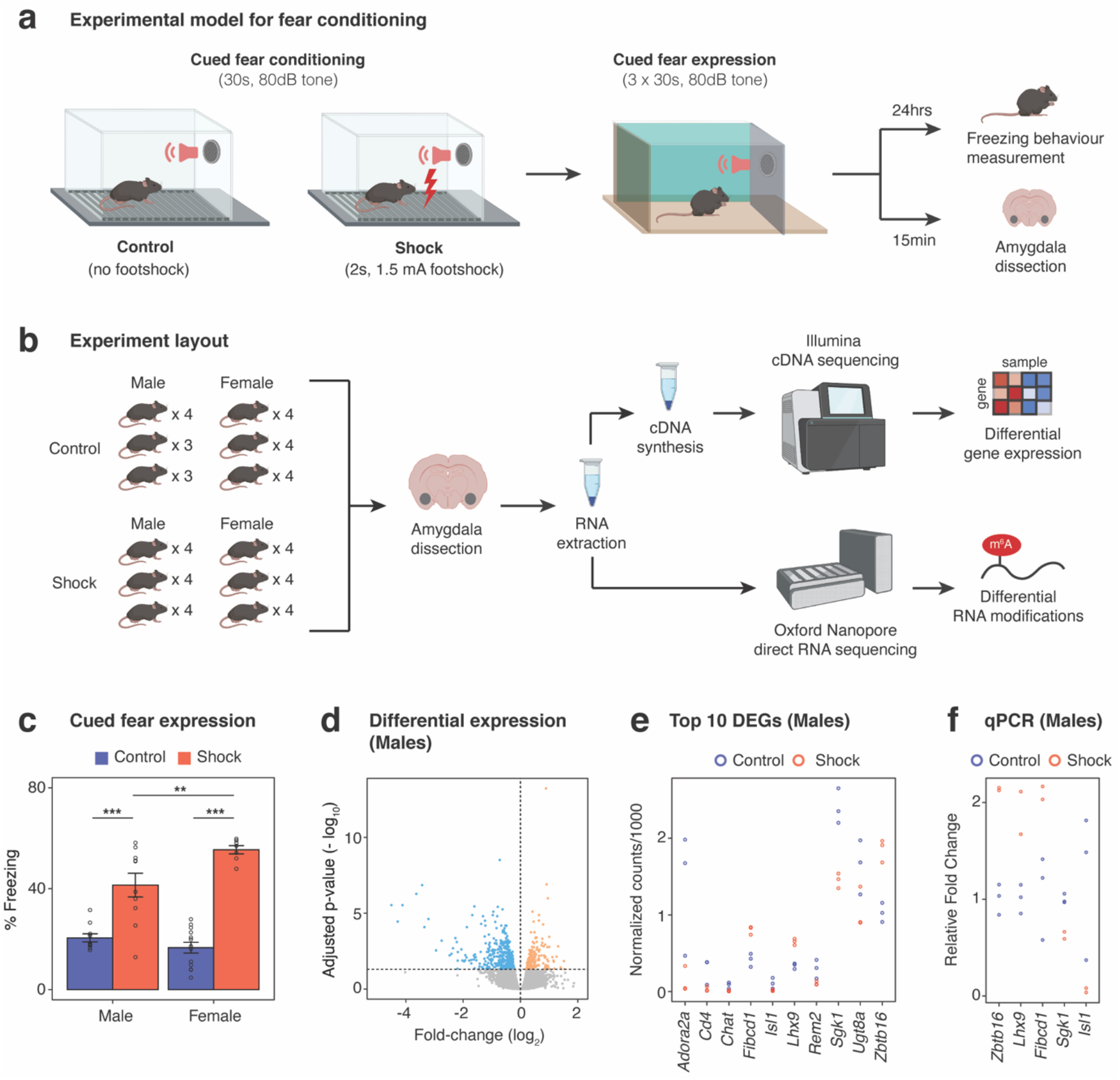
Sex-specific transcriptional profiles are associated with the acquisition of PTSD-like fear memories in the amygdala. (**a**) Experimental workflow. (**b**) Adult male and female mice underwent cued-fear conditioning (or the control condition) after which they were tested for cued-fear expression 24hrs later; another group was sacrificed within 15mins of fear conditioning to obtain amygdala RNA which was subsequently pooled for indirect (Illumina) and direct (Oxford Nanopore) sequencing. (**c**) Face validity of the cued fear conditioning paradigm was verified in male and female mice which showed strong expression of conditioned fear responses. ***p<0.001 relative to non-fear conditioned controls, **p<0.01 for male vs female conditioned mice. n=7-10/group. (**d**) Differential gene expression in the amygdala of male mice; black dots in blue shade represent significantly downregulated genes and black dots in pink shade represent significantly upregulated genes. Thresholds for DESeq2: |FC|>20% and FDR <0.05. (**e**) Top 10 differentially expressed genes in male conditioned mice. (**f**) qPCR validation for selected differentially expressed genes in males.

These findings led us to investigate whether this significant sex-difference in expression of PTSD-like fear memory could be associated with differences in transcriptional and epitranscriptional processes associated with the initial acquisition of the fear memory. To address this, a separate group of mice was sacrificed within 15 min of the fear conditioning procedure, capturing the earliest molecular signatures of fear acquisition. Messenger RNA (mRNA) was extracted and isolated from the amygdala of each individual, pooled in groups of 3-4 individuals, and analysed by short-read RNA-seq to assess differential gene expression (DGE) responses to the PTSD-like stressor (**Figure 1a, Supplementary Tables 1-3**). We obtained on average 37.9 million read pairs per sample (SD = 7.23 × 10^6^; **Supplementary Table 1**). In each library, most of the reads were successfully mapped to the reference genome (84.72% ± 0.56) and subsequently assigned to an annotated gene (65.83% ± 1.03; **Supplementary Table 1**).

We identified 190 up-regulated and 401 down-regulated genes in conditioned male mice (fold-change > 0.2 and FDR < 0.05; **Figure 1d, Supplementary Table 2**). The expression of the top 10 DGEs in males included Zinc finger and BTB domain containing 16 (*Zbtb16*), LIM homeobox protein 9 (*Lhx9*), fibrinogen C domain containing 1 (*Fibcd1*), serum/glucocorticoid regulated kinase 1 (*Sgk1*), and Isl1 transcription factor, LIM/homeodomain (*Isl1*), all of which were qPCR validated using the same samples that were sequenced (**Figure 1e, f**).

*Zbtb16* was found to be significantly upregulated in conditioned male mice and is a transcriptional regulator involved in a myriad of processes including those linked to neurodevelopment^18, 19^. *Zbtb16* knockout mice show several behavioural impairments relevant to neurodevelopmental disorders such as autism spectrum disorder and schizophrenia. In addition to impaired cortical morphology, these mice also show dysregulation in genes associated with myelination and neurogenesis^20^. Functional expression of this gene in the adult hypothalamus has been recently linked to regulation of metabolism^21^. *Lhx9* and *Isl1* are transcription factors which also have roles in neurodevelopment^22, 23^. *Fibcd1* belongs to a class of proteins known as fibrinogen-related domains (FreD) with multiple functions which include innate immunity and are expressed in glial cells in the human brain^24^. To our knowledge the functional expression of these four genes in the adult amygdala and in response to PTSD paradigms such as the one used in our study has heretofore not been characterised. They therefore represent interesting targets for future investigations.

*Sgk1* was downregulated in expression. It is involved in several functions which include regulating ion channel activity, interfering with transcription and neuronal excitability^25^. In the amygdala, an earlier microarray study reported an increase in *Sgk1* expression following cued-fear conditioning which contrasts with our findings; however, these experiments were carried out using a different conditioning protocol and analysis of amygdala RNA was at later time points^26^. It is worth noting that *Sgk1* is also expressed in other brain regions including the hippocampus, where it has a role in contextual fear memory formation^27^, and in the prefrontal cortex (PFC). A downregulation in expression has been reported in post-mortem tissue analyses of PFC tissue from patients who had suffered from PTSD, and is associated with increased expression of contextual fear memory in rats^28^.

Independent Gene Set Enrichment Analysis (GSEA) for conditioned male mice indicated a significant over-representation of genes implicated in synaptic activity (**Supplementary Table 3**), a process that is well-known to be implicated in the acquisition of memories^29-32^.

To our surprise, despite the increased expression of cued-fear memory, fear conditioning induced a modest transcriptional response in conditioned female mice (**Supplementary Table 4**). We observed a significant decrease in expression for only a single gene, *Rps27A*; however, this was not validated using qPCR. There remains debate as to whether the amygdala is a sexually dimorphic structure^33^. Evidence suggests there are intrinsic differences linked to sex hormones that may explain our sex-specific observations in the formation and expression of emotional memories^34-37^. Our findings contrast with several studies which have examined sex-differences in PTSD-like phenotypes, albeit in rats, with different inducing paradigms. Acute predator scent stress (PSS) induced significant differential expression of amygdala genes in males and female rats, detected using a microarray screen, that were classified as having an extreme behavioural response^38^. A subsequent re-analysis of this data revealed sex-differences in cellular composition of the amygdala but no effect of PSS. Controlling for cell subtype revealed shared as well as dimorphic gene expression profiles and enrichment analyses in the amygdala associated with EBR^39^. Using a single-prolonged stress (SPS) which consisted of 2h restraint stress, 20min forced swim followed by brief exposure to diethyl ether or exposure to a live predator, male rats exhibited a significantly higher acoustic startle, and increased sensitivity to dexamethasone-induced suppression of the HPA-axis response compared to females. SPS decreased social interaction and sucrose preference females^40^. Gonadal hormone status of males had no effect on SPS-induced startle hyperreactivity or dexamethasone responses but did influence female sucrose preference. Analysis of the right basolateral amygdala revealed a paradoxical increase in *c-fos* expression only in females^41^. It is worth noting that for all the above studies, the transcriptional analyses was done on brain tissue samples collected from 8 days to 2 weeks after the exposure to the traumatic stimuli. There have been several other studies which have examined changes to the transcriptional architecture in the brain in female rodents following chronic adult and early life stress exposure^42-45^. Our understanding of the immediate changes to the transcriptional architecture in the brain in response to stress in female rodents and how this sets the stage for longer term sequelae is still limited. Overall, our DGE data suggests significant distinctions exist during the early transcriptional events of fear acquisition in males and females warranting much further investigation.

Recent evidence suggests that epitranscriptomic modifications, such as N6-methyladenosine (m6A), may have widespread regulatory roles in the brain^5-8^. ONT dRNA sequencing has the potential to enable comprehensive profiling of the diverse array of RNA base modifications that together comprise the ‘epitranscriptome’^46, 47^. However, this is a nascent field and analytical best practices are yet to be established^48^.

To explore epitranscriptome dynamics during fear acquisition, we performed dRNA sequencing on the same amygdala RNA samples as described above. Each sample was sequenced on a single PromethION flow cell, yielding a median of 3.9 million QC-passed reads per sample, each of which represents a native mRNA transcript (**Supplementary Table 5**). This enabled a broad survey of expressed transcripts (Average read count > 3), with 12,792/32,604 (39.2%%) of protein coding genes reaching an average read count greater than 15, the minimum threshold required for detection of putative RNA modifications (**Supplementary Table 6**).

An overview of our analysis workflow used to profile RNA modifications is shown in **Figure 2a**, exemplified using the transcript for Neurensin-2 (*Nrsn2*), a known modulator of emotional behaviour and putative biomarker for PTSD^49, 50^. Briefly, we used a combination of two recently developed software packages, *Xpore*^51^ and *Nanocompore*^52^, to detect differential RNA modifications in the comparison of conditioned vs controls samples. *Xpore* identified 50,681 candidate modification sites across 9,902 transcripts for males and 37,998 candidates across 8,979 transcripts for females (**Figure 2a,b**). *Nanocompore* identified 43,172 candidate modification sites across 1,547 transcripts for males and 33,216 candidates across 1,482 genes for females (**Figure 2a,b**). The tools use alternative metrics and statistical approaches to identify modified k-mers and concordant results provide the best evidence of modified sites. To maximise stringency, we retained only those candidates that were identified by both *Nanocompore* and *Xpore* and collapsed any candidates within 10 nt into a single modification site (**Figure 2a**). For males, this approach identified 7,487 candidate modifications between conditioned vs control conditions that were collapsed into 3,397 unique differentially modified sites (**Figure 2b**). In females, 4,808 candidates were collapsed into 2,331 unique differentially modified sites (**Figure 2b**). These high confidence sites showed mean ∼1.7-fold higher modification rates, as measured by displacement of the current signal, under the stress condition in both males (stress = 0.48, control = 0.27) and females (stress = 0.49, control = 0.29; **Figure 2c**;).

**Figure 2.**
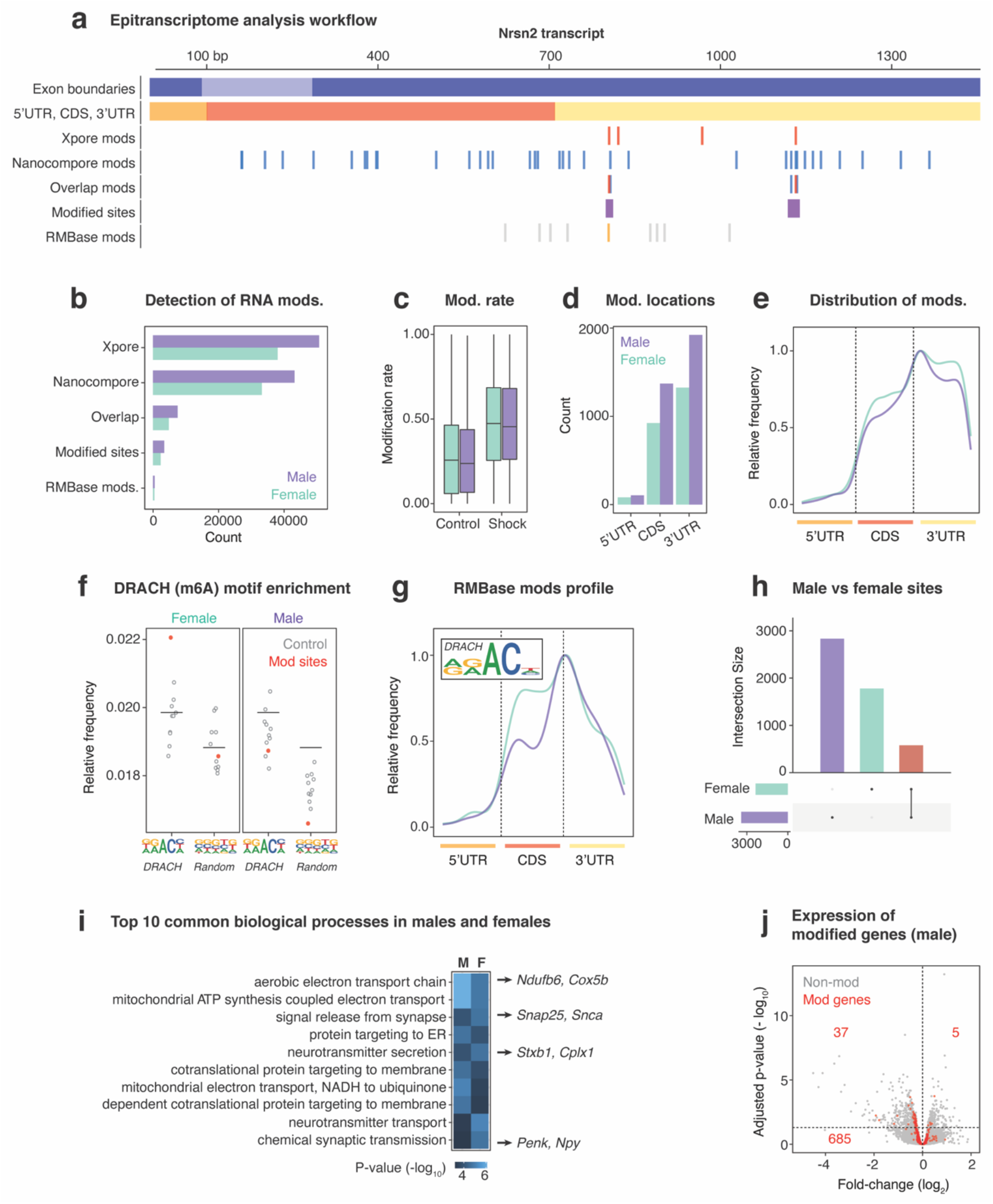
Epitranscriptome signatures of PTSD-like fear acquisition in the amygdala. (**a**) Schematic example using the *Nrsn2* mRNA transcript to illustrate the analytic workflow for profiling RNA modifications in conditioned vs control mice. Briefly, candidate modifications were identified separately by *Xpore* and *Nanocompore*. Only overlapping/adjacent candidates identified by both tools were retained and multiple candidates within 10 nt were collapsed into high confidence modified sites. Finally, modified sites were cross-checked against RMBase. (**b**) The number of candidates and modified sites retained at each step of the workflow just described (**c**) The modification frequencies recorded by *Xpore* for modified sites in stress vs control and male vs female samples. (**d**) The total number of modified sites identified within different regions (5’UTR, CDS and 3’UTR) of mRNA transcripts. (**e**) The positional distribution of modified sites, averaged across the whole transcriptome, within the same regions. (**f**) The relative frequency of canonical DRACH motif Kmers vs random k-mers observed within modified sites (blue), as opposed to random sites in matched transcript regions. (**g**) The positional distribution and k-mer frequency profile of modified sites that also overlapped RMBase modifications, with both showing characteristic features of m6A. (**h**) The number of modified sites identified that were sex-specific or shared between males and females. (**i**) The top ten common GO terms enriched among differentially modified genes in males and females. Example genes underlying each term are highlighted. (**j)** Volcano plot showing gene expression fold-changes observed between stress vs control, with genes that were identified as differentially modified highlighted in red. Minimal overlap between differentially expressed and differentially modified genes is apparent.

Most high-confidence modified sites were found within 3’ untranslated regions (UTR) of mRNA transcripts, with an enrichment close to the boundary with the coding sequence (CDS) observed in both males and females (**Figure 2d,e**). A smaller proportion of modified sites were found within 5’UTRs, which are generally shorter than 3’UTRs, but 5’UTRs exhibited the highest density of sites per nucleotide compared to the 3’UTR and CDS in both sexes (**Supplementary Figure 1a,b**). These observations are consistent with positional distributions previously reported for m6A and other known RNA modifications^53-55^.

Although *Nanocompore* and *Xpore* can identify differentially modified bases, neither tool determines the specific type of RNA modification at a given site. To further characterise and validate our modification sites, we assessed the enrichment of sequence k-mers consistent with the canonical m6A DRACH motif^56^. We found an enrichment of DRACH motif k-mers within high-confidence modification sites, compared to 10 sets of random non-modified sites, and no enrichment for k-mers of a randomly selected motif (**Figure 2f**). We also intersected our set of high-confidence modification sites with RMBase; a database of known RNA modifications^57^. We found 506 annotated RMBase modifications overlapping differentially modified sites in males and 421 in females, with almost all of them identified as m6A (**Figure 2b**). We retrieved the consensus sequence around these m6A sites and consistently observed the DRACH motif for both sexes (**Figure 2g**). Concordance with previously annotated RNA modifications, and adherence to known patterns of transcript position and sequence context confirms the reliability of our catalogue of differentially modified sites during fear acquisition.

Next, we functionally characterised genes and transcripts identified as being differentially modified. Out of 3,397 and 2,331 modified sites detected in males and females (**Supplementary Tables 7, 8**), respectively, we found 580 common modifications in 186 genes (average = 3.11 sites/gene; **Figure 2h**; **Supplementary Figure 1c**), suggesting a degree of both shared and sexually dimorphic epitranscriptome dynamics. GSEA for males and females indicated enrichment of equivalent and/or related gene ontology terms (**Figure 2i**). Within the top 20 most significantly enriched terms, 60% were found in both males and females (**Supplementary Tables 9,10**).

Mitochondrial function and neurotransmitter production/secretion were the predominant functional categories identified by GSEA (**Figure 2i**). The former is particularly interesting in the context of evidence implicating mitochondrial function in the stress response^58-60^ and mitochondrial dysfunction in PTSD pathology^61, 62^. Mitochondrial metabolic processes are components of the cell danger response (CDR) which is activated by stressors. Negative feedback mechanisms exist to ‘switch off’ the CDR once the stress has passed but if they fail, this may contribute to oxidative stress and inflammation^63^.

Among the top differentially modified gene candidates were multiple synaptic genes previously implicated in PTSD, including Preproenkaphelin (*Penk*)^64^ and Neuropeptide Y (*Npy*)^65^. Evidence suggests the *Npy* system promotes resilience-to-stress in rodents and reduced NPY in humans is associated with PTSD^65^. We observed differentially modified sites in the 3’UTR and CDS regions of *Npy* in males and females. In each case, the modification frequency was upregulated during fear conditioning, whereas expression of *Npy* mRNA was not found to be upregulated, suggesting transcription-independent regulation of *Npy* by RNA modifications may occur during fear conditioning. The lack of concordance between differential modification and expression in *Npy* was similarly observed across the transcriptome, with just 150/3,397 (4.41%) of high confidence modifications being harboured within differentially expressed genes (**Figure 2j; Supplementary Table 11**). This finding reveals widespread expression-independent epitranscriptomic regulation in the amygdala during fear acquisition and is consistent with previous work showing an absence of correlation between changes in gene expression, translational efficiency, and m6A-methylation^66^. We hypothesise that dynamic RNA modification processes are among the earliest molecular events in fear conditioning and may regulate the activity of key effector genes prior to activation of mechanisms which include but are not limited to differential gene expression in a sex-specific manner.

Gaining an insight into the molecular mechanisms that are triggered immediately following trauma are important to comprehend how they are implicated in the neuropathological cascades that culminate in the onset of PTSD. Our study provides a framework for exploring acquisition-induced transcriptomic and epitranscriptomic events underpinning PTSD-like phenotypes in male and female mice and implicates RNA modifications in the regulation of fear. There is broad scope for future work, but one major issue must be addressed; how does one interrogate the functional roles of RNA modifications^67^?

For some modifications such as m6A, the majority of known m6A sites are unmethylated at baseline^68^. The functional relevance of constitutive versus regulated m6A sites is unknown but such sub-stoichiometric levels would indicate a large margin in which to regulate RNA metabolism and gene expression. Furthermore, while the lifecycle of modifications is thought to be transient, this may vary from transcript to transcript. Our data are also consistent with recent work showing that RNA modifications can form clusters on the same transcript^69^, exerting cumulative or combinatorial effects on RNA metabolism. One low resolution approach to providing answers is by manipulating the expression and activity of epitranscriptomic machinery enzymes - ‘writers’, ‘erasers’ and ‘readers’. This has been used by numerous studies in the context of learning, memory, and the stress response^66, 70, 71^. There have also been several pharmacological compounds developed to inhibit or activate these enzymes^72, 73^. One higher resolution strategy involves the use of CRISPR-Cas Inspired RNA-targeting (CIRT) system^74, 75^ but this is yet to validated in vivo. And lastly, future investigations should consider incorporating techniques such as ribosome profiling and proteomics^76^.

In conclusion, our study opens a new and exciting frontier in molecular psychiatry research that has potential to reshape our understanding of stress-induced psychiatric disorders. RNA-based therapies have found success in treatment of diseases from cancer to COVID19^77^. Perhaps, there is scope to treat psychiatric disorders such as PTSD in a similar manner.

## Acknowledgements

AG wishes to thank Dr Nicholas Everett and Dr Erin Lynch for technical assistance and the Laboratory Animal Services team at the Brain & Mind Centre for assistance with animal husbandry. We thank our colleagues Derrick Lin and Warren Kaplan for providing technical support and oversight of high-performance computing at the Garvan Institute. AG also wishes to thank Professor Tim Bredy (University of Queensland) for the many intellectually challenging conversations associated with this study.

## Availability of data and materials

All data generated or analysed during this study are included in this published article and its supplementary information files. Sequencing files have been uploaded to NCBI GEO (PRJNA779821). Scripts for analysis of data are available on request.

## Author contributions

AG, AR, JCA and ID designed the study. JCA advised on the fear conditioning experiments. AG managed the project, carried out all stress procedures, behavioural experiments, dissections and RNA extractions. AR, JH and IS carried out sequencing of samples. AG and AR analysed the data and wrote initial drafts of the manuscript. JCA, ISM, ID provided advice on data analysis and interpretation. All authors contributed to the interpretation of the data, critically revised the manuscript, read and approved the final version before submission.

## Ethics approval and consent to participate

All experiments were approved by the University of Sydney Animal Ethics Committee (2018/1425). All efforts were made to minimize animal suffering and to reduce the number of animals used.

## Supplementary Material

### Materials & Methods

#### Animals & Housing

All experiments were conducted in accordance with the NHMRC Australian code for the care and use of animals for scientific purposes and approved by the Animal Ethics Committee of University of Sydney (AEC2018/1425). All efforts were made to minimise animal suffering and to reduce the number of animals used. Adult male and randomly cycling female C57BL/6J mice were obtained from the Animal Resources Centre (Western Australia) and housed in groups of 5 in individually ventilated cages (IVC) at the Brain & Mind Centre (New South Wales) in a temperature/humidity-controlled environment (21°C, 55.5%). Following 1 week of acclimatisation to reverse-phase lighting (lights ON: 21h00, OFF: 09h00), mice were singly housed in IVCs for an additional week. Food and water were available *ad libitum*.

#### Cued Fear Conditioning & Cued Fear Memory Testing

The procedure for conditioning and testing of cued-fear memory is based on recently published work ^1-3^. Briefly, mice were fear conditioned in a wooden sound-proof chamber, transparent front and rear-facing walls, opaque side walls with a white light and a metal grid floor (Context A). The conditioning session started with a 198 s acclimation period which was followed by a 30 s 80dB tone (9KHz sinewave, conditioned stimulus, CS) that co-terminated with a 2s 1.5mA foot-shock delivered through the metal grid as a constant current (unconditioned stimulus, US). The conditioning session ended with a 60 s after-shock interval. Non-shock control mice were exposed only to the CS tone. Fear conditioning chambers were cleaned using 70% ethanol spray solution. Within 15 mins of being fear conditioned, one group of stress and control mice were culled, brains extracted and snap frozen. Cued fear memory was tested in another group of stress and control mice 24hrs later in a novel context (Context B: wooden sound-proof chamber, red light, red perspex flooring, peppermint oil scent). Following a 2 min baseline period, all mice were presented with 3 × 30 sec CS with an inter-tone interval of 1 min. The memory retrieval session ended with a 60 s interval after the last tone was presented. Chambers were cleaned using F10 disinfectant spray solutions. Freezing was measured in response to the CS using the CleverSys FreezeScan® video tracking system and software.

#### Tissue Collection & RNA Extraction

Mice were sacrificed by cervical decapitation within 15 min of being fear conditioned. Brains were extracted, snap-frozen in liquid nitrogen and stored at -80°C. As previously described ^4^, bilateral punches (0.5 to 1mm) of the amygdala (AMG: central and basolateral amygdala) were made using the Paxinos and Watson Atlas as a guide ^5^. Tissue was collected in RNA-free tubes and RNA was extracted using Qiagen RNeasy™ Micro kit (Qiagen, MD, USA) according to the manufacturer’s instructions. RNA concentrations were quantified using a NanoDrop™ One spectrophotometer (Thermofisher Scientific®, MA, USA) and only samples with 260/280 ratios of greater than 1.7 were used for downstream analyses. Samples from 3-4 mice were randomly pooled for subsequent analyses by Illumina NextSeq and Promethion. mRNA was purified from total RNA pools using the Dynabeads® mRNA Purification Kit (Thermofisher Scientific®, MA, USA) according to the manufacturer’s instructions.

#### Illumina RNA sequencing

Sequin RNA Mix (Hardwick et al. 2016) sequencing control was added at a fraction of 1% of total purified mRNA. Purified mRNA with Sequin was used to prepare libraries using the KAPA Stranded RNA-Seq Library Preparation Kit (Roche, CA, USA) with SeqCap Adapters (Roche CA, USA) as per manufacturer’s instructions. The libraries were sequenced on an Illumina NextSeq 500 System, generating 24.6-48.9 million, 2 × 150bp read-pairs per sample.

#### Nanopore direct-RNA sequencing

Direct RNA sequencing was performed using the ONT Direct RNA Sequencing Kit (SQK-RNA002), as per the manufacturer’s instructions. ∼200 ng of purified mRNA per sample was provided as input and optional first-strand cDNA synthesis was performed using SuperScript III Reverse Transcriptase (Thermo Fisher Scientific). For each sample, ∼30-50 ng of prepped library was loaded onto a single ONT PromethION flow cell and sequenced for a maximum ∼72 hours, generating 2.2-4.6 million native RNA reads per sample.

#### Bioinformatics – Gene expression profiling

Reads were trimmed using Trim Galore (Galaxy version 0.6.3) and subsequently aligned to the reference Mus musculus genome (GRCm38) using HISAT2 (Galaxy version 2.1.0). Reads were annotated (Gencode v.M25) and counted using featureCounts (Galaxy version 2.0.1). Biostatistics and visualization were run in R (version 3.6.3) with the Rstudio GUI (version 1.2.5033). For each comparison – male: stress(3) v control(3), female:stress(3) v control(3) - genes were filtered to have at least 5 reads in at least 2 samples for each gene and have gene biotype of protein-coding, long non-coding or microRNA. RUVSeq ^6^ and DESeq2 ^7^ were then used to quantify differential gene expression between groups. We set a threshold criteria of a minimum 0.2 expression fold-change and DGE FDR<0.05. Gene ontology and enrichment analysis was performed using EnrichR with a significance threshold set at FDR <0.05 ^8^.

#### qPCR Validation

RNA was reverse-transcribed to complementary DNA using the Applied Biosystem® High Capacity cDNA Reverse Transcription Kit (10X RT Buffer, 25X dNTP mix (100mM), 10X RT Random Primers, Multiscribe® Reverse Transcriptase) on the Applied Biosystem® GeneAmp PCR System 9700 (Thermofisher®, Waltham, MA, USA). qRT-PCR was carried out on the StepOnePlus® PCR machine (Thermofisher®, Waltham, MA, USA) using the following primer assays designed by Integrated DNA Techologies (Skokie, Illinois, USA). Samples were heated to 95°C x 10 min, and then subjected to 40 cycles of amplification by melting at 95°C x 15 s and annealing at 60°C x 1 min. Experimental samples were run in duplicates with 1.33 μL complementary DNA (cDNA) per reaction. To check for amplicon contamination, each run also contained template free controls for each probe used. PCR data were normalized using β-actin and transformed using the ΔΔCt method.

#### Bioinformatics – Epitranscriptome profiling

To generate a comprehensive catalogue of high-confidence RNA modification sites present in the amygdala of male and female mice exposed to conditioned fear, we used two recently released software Nanocompore and Xpore on our direct RNA-seq libraries. Nanocompore collects current intensity and dwell time at each position, as the native RNA goes through the pore, and uses those variables to perform pairwise statistical comparisons between control and stress samples. The tool reports positions with a significant statistical difference for each of the individual variables, but also models a logistic regression that takes both current intensity and dwell time into account in search for significant differences. To minimise the false-positive discovery rate, we applied strict filtering to the results, only retaining putative modifications with a significant p-value (alpha=0.01) in all three statistical tests. Xpore, on the other hand, models the current intensity as a mixture of two Gaussian distributions, for modified and unmodified RNA, using prior information to help guide parameter estimation. Then given two conditions, such as control and stress, the tool is able to identify differential modification, but also quantitate them. We only selected putative modifications with significant differences in modification rates (t-test, p-value < 0.01) between the conditions and that were increasingly modified in the stress mice. To further increase the confidence in our dataset, we only retained modifications identified by both Nanocompore and Xpore, and that were at most 10 nt apart from each other and we also collapsed modifications less than 10 nt apart into a single modification site. We compared the coordinates of the modification sites identified using this approach against modification sites previously annotated in RMBase v2.0. We also performed gene set enrichment analysis using EnrichR with a significance threshold set at FDR <0.05.

---

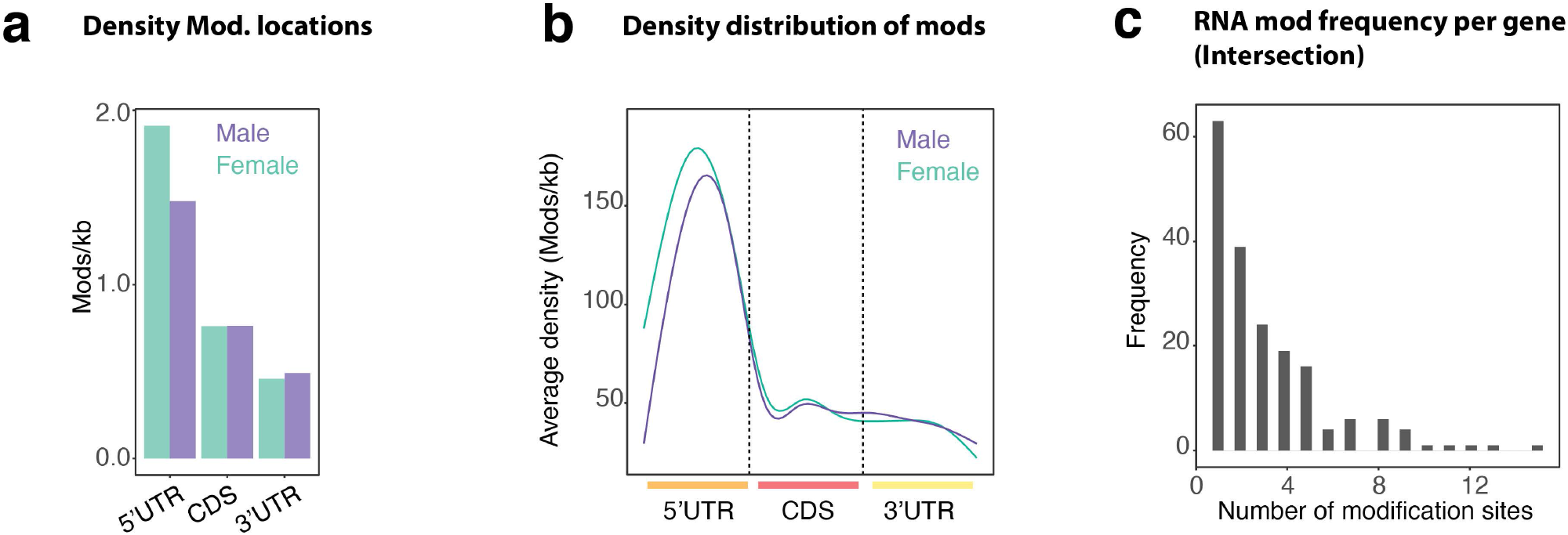

